# An epigenomic shift in amygdala marks the transition to maternal behaviors in alloparenting virgin female mice

**DOI:** 10.1101/2021.05.14.444198

**Authors:** Christopher H. Seward, Michael C. Saul, Joseph M. Troy, Payam Dibaeinia, Huimin Zhang, Saurabh Sinha, Lisa J. Stubbs

## Abstract

In many species, adults care for young offspring that are not their own, a phenomenon called alloparenting. However, most nonparental adults must be sensitized by repeated or extended exposures to newborns before they will robustly display parental-like behaviors. To capture neurogenomic events underlying the transition to active parental caring behaviors, we analyzed brain gene expression and chromatin profiles of virgin female mice co-housed with mothers during pregnancy and after birth. After an initial display of antagonistic behaviors and a surge of defense-related gene expression, we observed a dramatic shift in the chromatin landscape specifically in amygdala of the pup-exposed virgin females, accompanied by a dampening of anxiety-related gene expression. This epigenetic shift coincided with hypothalamic expression of the oxytocin gene and the emergence of behaviors and gene expression patterns classically associated with maternal care. The results outline a neurogenomic program associated with dramatic behavioral changes and suggest molecular networks relevant to human postpartum mental health.

## INTRODUCTION

Interactions between newborn animals and their parents are profoundly important, being critical to the well-being of the offspring and intensely consequential to the parents as well. In most mammals, parental care is typically relegated to the female that bears the offspring, with hormonal shifts that occur during pregnancy and the early postpartum period priming her for this experience. These dramatic hormonal shifts also alter a mother’s morphology, physiology, and brain structure in ways that persist far beyond the initial parenting experience [1,2]. In addition to these physical changes [3], mothering also alters a female’s behavior, in both the immediate and the longer-term. In particular, the sight, sounds, and odors of newborns – which may otherwise be perceived as aversive by adults – become intensely rewarding and motivating to the mother [4–6]. As with other changes associated with parenting, the shift from aversion to intense affiliation and reward is coordinated by steroid hormones and a rapid surge in neuropeptide secretion around the time of birth [3]. Most significantly, a surge of oxytocin, stored during pregnancy within the paraventricular and supraoptic nuclei of the hypothalamus [7], is released to target neurons within a brain circuit central to fear, aversion, reward, and the evaluation of emotional salience [8].

New mothers are not the only individuals that can experience this switch to pup-affiliative behaviors. For example, although virgin female rats display a clearly aversive response to pup stimuli, this response can be overcome by the process of sensitization, which involves a series of repeated interactions; after sensitization, virgin rats will display robust maternal behaviors – hereafter referred to as maternal behaviors for simplicity – with pups [6]. In contrast, adult virgin female mice do not display any obvious sign of aversion and will instead spontaneously display certain maternal behaviors shortly after given first access to young pups [9,10]. Sensitization enhances this response; virgin female mice repeatedly exposed to pups significantly increases both the range and intensity of maternal behaviors [11]. Intriguingly, it has been shown virgin female mice continuously co-housed with new mothers will display maternal behaviors more rapidly under the instruction and encouragement of the mothers, a process that depends upon the activation of oxytocin neurons [12]. Like mothering itself, this experience of caring for young that are not one’s own, or alloparenting, impacts future behavior. For example, juvenile female rats that have had the experience of “babysitting” younger siblings are highly motivated to display maternal behaviors in future encounters with pups [13], and sensitized adult virgin female mice demonstrate enhanced parenting skills when they have their offspring of their own [11,14,15]. Indeed, many of the mechanisms that reshape a mother’s brain and behavior also appear to operate in alloparenting females, where intriguingly, they are activated without the hormonal priming stimulated by pregnancy, parturition, and nursing.

Here, we investigated the functional genomics profile of the brains of co-housed virgin female mice as they transitioned from pup-naïve to a robust display of alloparenting behaviors toward pups. To identify genes modulated during this transition, we examined alterations in gene expression in multiple brain regions over several days of continuous pup exposure. Because histone modifications have been implicated as central to the behavior of both new mothers and sensitized virgins [16], we also investigated chromatin accessibility profiles using H3K27Ac (histone H3 acetylated at lysine 27), a marker of open chromatin, in the same brain regions. The data reveal defense-related neurogenomic pathways that are silenced, and others that are activated, across the brains of co-housed alloparenting virgins as they transition to maternal behaviors and confirm an active role for chromatin remodeling in this behavioral switch, especially within the amygdala.

## RESULTS

### Antagonistic behavior, followed by active nurturance in virgins co-housed with mothers and pups

In the most common version of rodent pup sensitization experiments, the pups are placed into the cage of a virgin female for short periods, then removed to be fed, repeatedly over the course of several consecutive days [11]. However, over years of mouse breeding, we had observed that nulliparous females co-housed with nursing mouse dams and their litters will display maternal behaviors toward the pups, suggesting a way to achieve more continuous, longer-term interaction. Indeed, a recent study has demonstrated that in this context, virgins respond more quickly to the pups as they are actively instructed and encouraged by the mothers [12]. This co-housing paradigm provided us with an excellent opportunity to measure the brain’s functional genomic response to pups in the virgin females over time as they transition to maternal care. To document the timing of this transition, we co-housed four pairs of virgins and pregnant dams and filmed activity in the cages from early pregnancy though the fourth postnatal day (**File S1**; Files S1 and S2 available at https://trackhub.pnri.org/stubbs/ucsc/public/allo.html). For purposes of this study, we were primarily interested in the interactions between virgins and mothers, on the one hand, and virgins and pups one the other. Therefore, as a primary indicator of these interactions, we scored the virgins for pup-grooming and mother-grooming behaviors during 5-minute intervals at the top of each hour; summing the scores in each cage over 6-hour periods coordinated with the light/dark cycle, providing a useful summary of the overall behavioral patterns (**Table S1**). Throughout, the two females were most often found together, interacting or resting in the shared nest, and grooming each other regularly while awake throughout the observation period. In contrast, although the virgins began to investigate the pups immediately after birth, they did not begin licking and grooming the pups consistently until around postnatal day 2 (P2), after which we increasingly observed the virgins engaged in pup licking/grooming behavior (**Fig. 1A; Table S1)**. To test the hypothesis that pup-focused grooming increased for the virgins while mother-focused grooming did not, we selected data binned for 6 hours around 12:00 (the beginning of the dark period during lights-out) (**Fig. 1B**). Pup-focused grooming bouts significantly differed across days P1-P4 (repeated measures ANOVA, F_3,9_ = 9.91, p = 0.003), increasing over time, while mother-focused grooming bouts did not significantly differ across days P1-P4 (F_3,9_ = 0.67, p = 0.59).

**Figure 1.**
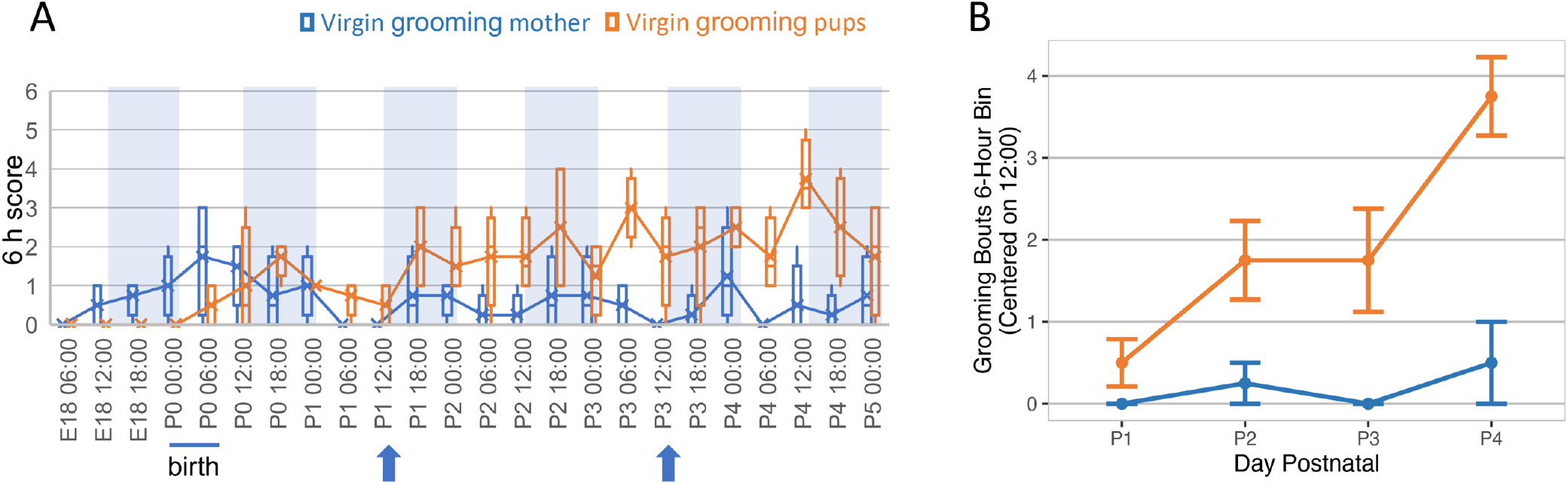
Behaviors exhibited by virgin females co-housed with new mothers and pups over six days beginning the day before birth. **(A)** Four cages of co-housed mothers and virgin females were recorded over a period of several days before and after birth and grooming behaviors (virgins to pups, plotted in orange; or virgins to mother, blue) were scored (0 or 1) in 5-minute intervals at the beginning of each hour from 12:00 am (0:00) the day before the birth (E18) through the end of the fourth postnatal day (P4), then scores were summed over 6 h periods. To generate the graph, 6 h summed scores were plotted for the four cages as box-and-whisker plots. Times shown mark the end of each 6h period scored. Blue shading in each plot shows the “lights out” periods (12 h beginning at 12:00 pm) for each day. All pups were born within a 6-hour period at the beginning of the light phase on the day designated as P0 for that particular cage, as marked with a bar below each graph. Behaviors plotted and colors used are shown above each graph. Blue arrows below each graph show the times of day that samples were taken from similarly co-housed pairs for gene expression and chromatin analysis. **(B)** The frequency with which co-housed virgins groomed pups (plotted in orange) increased significantly over days (*p*=0.003), as illustrated by a plot focused on 12:00 pm (start of lights out), while the frequency with which virgins groomed mothers (blue) did not change. Values plotted are mean ± standard error.

We also observed additional behaviors that are worth noting here. For example, as described by Carcea and colleagues [12], we observed mothers steering virgin cagemates that had wandered off to feed or explore back to the nest; often this involved the mothers grabbing the virgins by the base of the tail and actively pushing them to the nest and pups. Afterward, the mother would herself typically leave the nest to feed, leaving the virgin to care for the pups. Not described in the published study but displayed by all virgins we recorded here, we also observed signs of early antagonism toward the pups. Specifically, during the first two postnatal days we observed the virgins grabbing pups in their mouths and actively tossing them or pushing them out of the shared nest **(Table S1, examples of both behaviors in File S2)**. By P3, this behavior was no longer observed, as the virgins spent more time in the nests, licking and grooming the pups with increasing frequency in classic hunched or prone nursing postures (**Table S1**), similar to behavior documented for sensitized female rats [17]. Together, these observations suggested that we could indeed capture the transition from the possibly antagonistic pup-naïve state to robust pup affiliation between postnatal days 1 and 3 in this continuous-exposure paradigm.

### Hormone- and neurotransmitter-related genes are dynamically expressed throughout the virgin brain during the first three days of pup interaction

To understand the functional genomic underpinnings of this behavioral transition, we collected RNA from the brains of five virgin females co-housed with a pregnant dam at each of three time points: before birth (2 hr into the dark period of embryonic day 18, or E18) and at the same time during postnatal day 1 (P1) and P3. We collected and sequenced RNA from four brain regions involved in pup response, aversion, affiliation, and reward: hypothalamus, amygdala, striatum, and frontal cortex.

At P1, we saw an intense transcriptomic response in amygdala with very little expression change in other brain regions; by P3, relatively large numbers of differentially expressed genes (DEGs) were detected in both amygdala and hypothalamus (**Fig. 2; Table S2**). The transcriptomic response did not correlate simply with expression of immediate early genes (IEGs) such as *Fos*, which is classically used to mark neuronal activity [18]. However, the IEG *Npas4*, which has been implicated specifically in social recognition [19] and reward-related behaviors [20], was upregulated at P3 in all brain regions tested (**Fig. 3**). Focusing first on the hypothalamus, genes encoding neuropeptide hormones oxytocin (*Oxt*) and prolactin (*Prl*) were first up-regulated at P3, when the virgins were beginning to consistently display maternal behaviors (**Fig. 1, Fig. 3**); these hormones are central to initiation of maternal response in both mothers and alloparenting virgins [12,21,22]. At P1, dopaminergic (DA) signaling components including *Drd1* were down-regulated along with related Gene Ontology (GO) and functional categories such as morphine addiction and behavioral despair. However, *Drd1* returned to pre-exposure levels in hypothalamus at P3, at the same time that genes related to the activity of dopaminergic neurons were significantly up-regulated.

**Figure 2.**
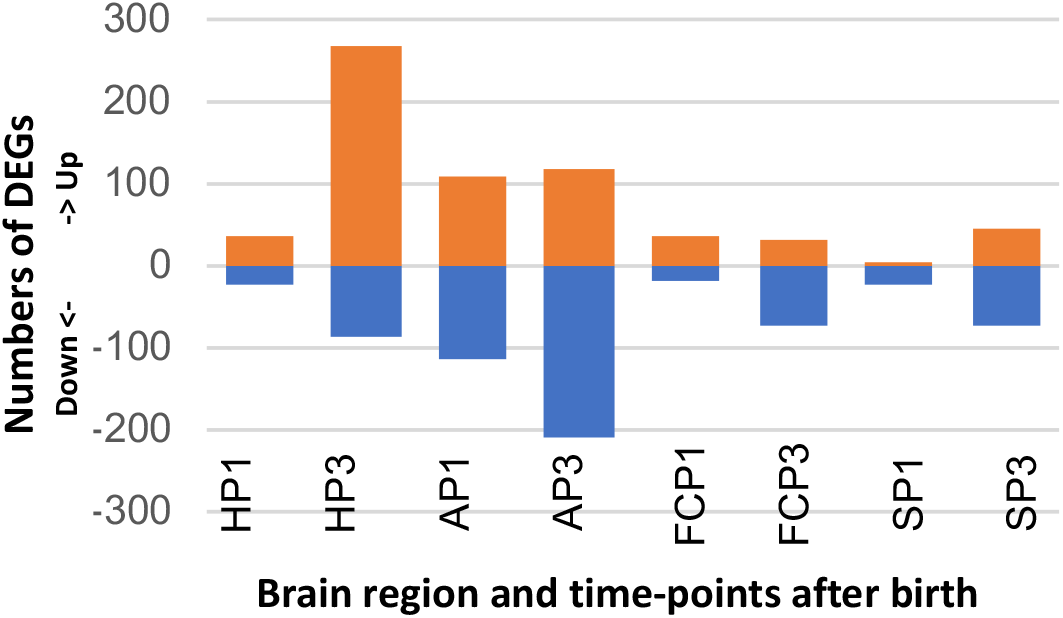
Numbers of genes up-(orange) or down (blue)-regulated in brains of pup-exposed compared to non-exposed virgin females over time. Numbers represent all genes identified as differentially expressed at fdr ≤ 0.05 in each set of pairwise comparisons. H=hypothalamus; A=amygdala; FC=frontal cortex; S=striatum; P1=postnatal day 1, P3=postnatal day 3.

**Figure 3.**
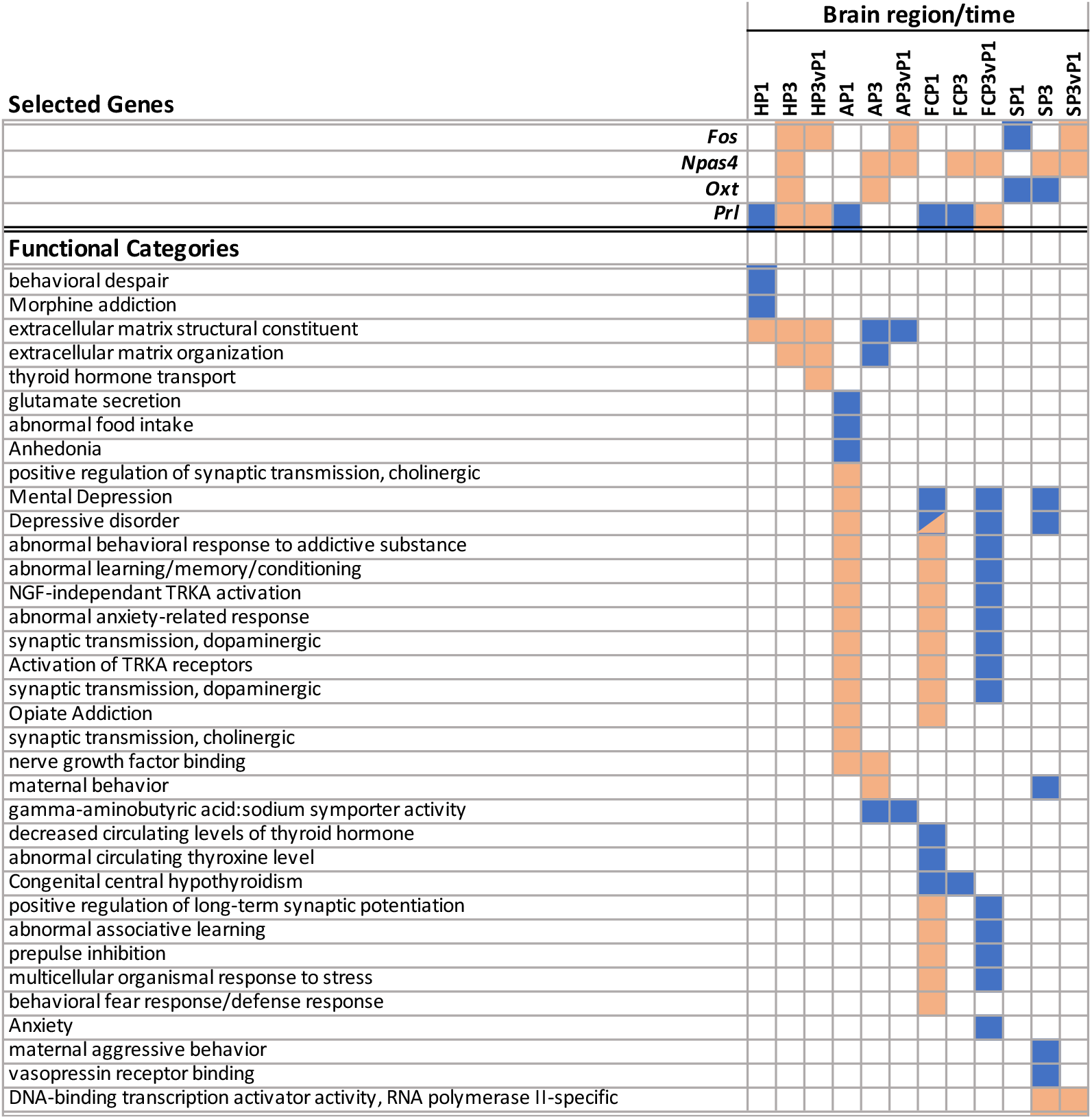
Enrichment of differentially expressed genes in functional categories. Differentially expressed genes (identified at fdr≤0.05 and with absolute value of fold change ≥1.5) were used to identify enriched functional categories using the ToppCluster tool (Kaimal et al., 2010), as described in Methods. Top panel shows differential expression levels for selected genes, as described in the text. Categories shown are a representative subset of the full report included in Table S3, with up- or down-regulation and category enrichment levels displayed as a heat map. Colored cells denote up-(orange) or down-regulation (blue) for Selected Genes (top panel) or Functional Categories (lower panel) in each brain region/time.

Therefore, a switch from repressed to increased dopamine-related gene expression was coordinated with the increase of *Oxt* and *Prl* expression in the hypothalamus. This pattern is similar to that observed in mothers at the time of birth and is consistent with the role of DA signaling in OXT and PRL release [23,24]. It is also consistent with recent observations from single-cell sequencing that show DA neurons in the hypothalamic preoptic area to be activated in maternally behaving animals [25]. The data suggested that a shift to a neuropeptide and neurotransmitter environment favoring stable maternal behavior was developing in the hypothalamus at P3, concomitant with increased expression of alloparenting behaviors in the virgin mice. Furthermore, in light of the hypothesized role of histone modifications on maternal behavior [16,26,27], it is also worth noting the low-level but coordinated up-regulation of genes encoding chromatin remodeling and binding proteins (*Hdac10*, *Hdac7*, *Sirt6*, and *Sirt7*, *Kmt5c*, *Smarcd3*, *Atrx*, *L3mbtl1*) which we observed in the virgin hypothalamus at P3. This coordinated shift suggested the existence of a subtle but significant epigenetic response in the hypothalamus around or before that time (**Table S2A**).

In striking contrast to hypothalamus, components of DA signaling were coordinately *up-regulated* in the amygdala at P1, along with genes encoding endogenous opioids, proenkephalin (*Penk*), and prodynorphin (*Pdyn*). The combined up-regulation of these genes led to P1 enrichment of multiple functional categories indicating that the virgin females were experiencing stress and anxiety during the first day after the birth of the pups (**Fig. 3; Table S3**). At P3, many of the anxiety-related amygdala DEGs had returned to pre-exposure levels or were down-regulated compared to E18 controls, suggesting that the initial P1 surge of transcription for these genes might be actively silenced. At the same time that the surge of anxiety-related genes was suppressed, the GO biological process category “maternal behavior” was identified as being enriched in P3 up-regulated genes (**Fig. 3**).

Frontal cortex tracked the amygdala closely in terms of DEGs, direction of change, and enriched functional categories, with a few notable exceptions. In particular, genes related to thyroid hormone activity were uniquely downregulated in the virgin frontal cortex at P3, a finding that is especially interesting given the known role of thyroid hormone in maternal care [28]. Furthermore, in addition to the dopamine-related genes similarly up-regulated in amygdala and cortex (**Table S2**), a second cadre of genes associated with depressive states, but related to abnormal thyroid hormone signaling, were down-regulated; the result was that depression-related functional categories were both up and down-regulated in the frontal cortex DEG set (**Fig. 3; Table S3**).

Finally, in P3 striatum, down-regulated categories were centered on neuropeptide-related genes including those encoding vasopression receptor (*Avpr1a*) and prolactin receptor (*Prlr*) (**Fig. 3**). On the other hand, the gene encoding neuropeptide cholecystokinin (*Cck*), which positively regulates striatal dopamine signaling in *Drd2*-expressing neurons [29,30] was up-regulated in striatum at P3 compared to P1. This event is notable, since *Cck* plays a critical role in the postnatal maintenance of maternal behaviors [31] and mediates responses to anxiety and reward [32,33]. Together these data indicated that a transcriptomic signature consistent with a “maternal response” - as it is classically defined by neuropeptide and neurotransmitter gene expression-was observed in the virgin mice beginning around P3. In particular, the P1 burst of anxiety-related genes was down-regulated to pre-exposure levels in amygdala and frontal cortex by this time. In contrast with this response in amygdala, DA signaling was *down-regulated* at P1, then *up-regulated* at P3 in the hypothalamus of the pup-exposed virgins, concordant with the onset of maternal behaviors in those mice.

### Comparison to published datasets

#### Parallels to gene expression in brains of new mothers

As referenced above, expression of several key markers that have been identified in new mothers was also observed in the alloparenting virgins at the P3 time point. An obvious next question was whether and how gene expression aligned more globally between maternal and alloparenting virgin brains. Most published maternal datasets were generated with distinct hypotheses and biological questions in mind, investigating brain regions and time points very different from ours, complicating direct comparisons. Nevertheless, two published data series warrant some discussion.

In the first series, Gammie and colleagues used microarrays to compare gene expression between virgins (not exposed to pups) and nursing females at P7, after maternal behaviors have been robustly established [34–37]. The same group later completed a meta-analysis of their data to identify genes that were commonly dysregulated across the maternal brain. Despite the differences in methods, time points selected, and brain regions examined, we noted that DEGs identified in the meta-analysis were enriched in similar GO categories, pathways, and disease associations to those we identified as most pronounced in the pup-exposed virgin brains: neuron development, addiction, mental health disorders, and pathways involving oxytocin, vasopressin, prolactin, and opioids [38]. The similarity suggests commonalities between maternal behavior and alloparenting behavior.

A second published data series examined maternal gene expression over a wide range of time points pre-and post-partum including P1 and P3, and importantly, used experimental and statistical methodology very similar to ours [39]. However, cortex (neocortex in the maternal study, which includes frontal cortex and additional cortical regions) and hypothalamus were the only brain regions examined commonly in both studies. This similarity allowed us to use formal statistical techniques to measure the degree of overlap between gene sets from our study and this previously collected dataset. Using a hypergeometric test to compare gene expression in pup-exposed virgins and mothers (**Table S2B, S2C**), we found that DEGs up-regulated in the hypothalamus of P3 virgins correlated positively and most significantly with genes up-regulated in the maternal hypothalamus at P10 (**Table 1**).

**Table 1.**
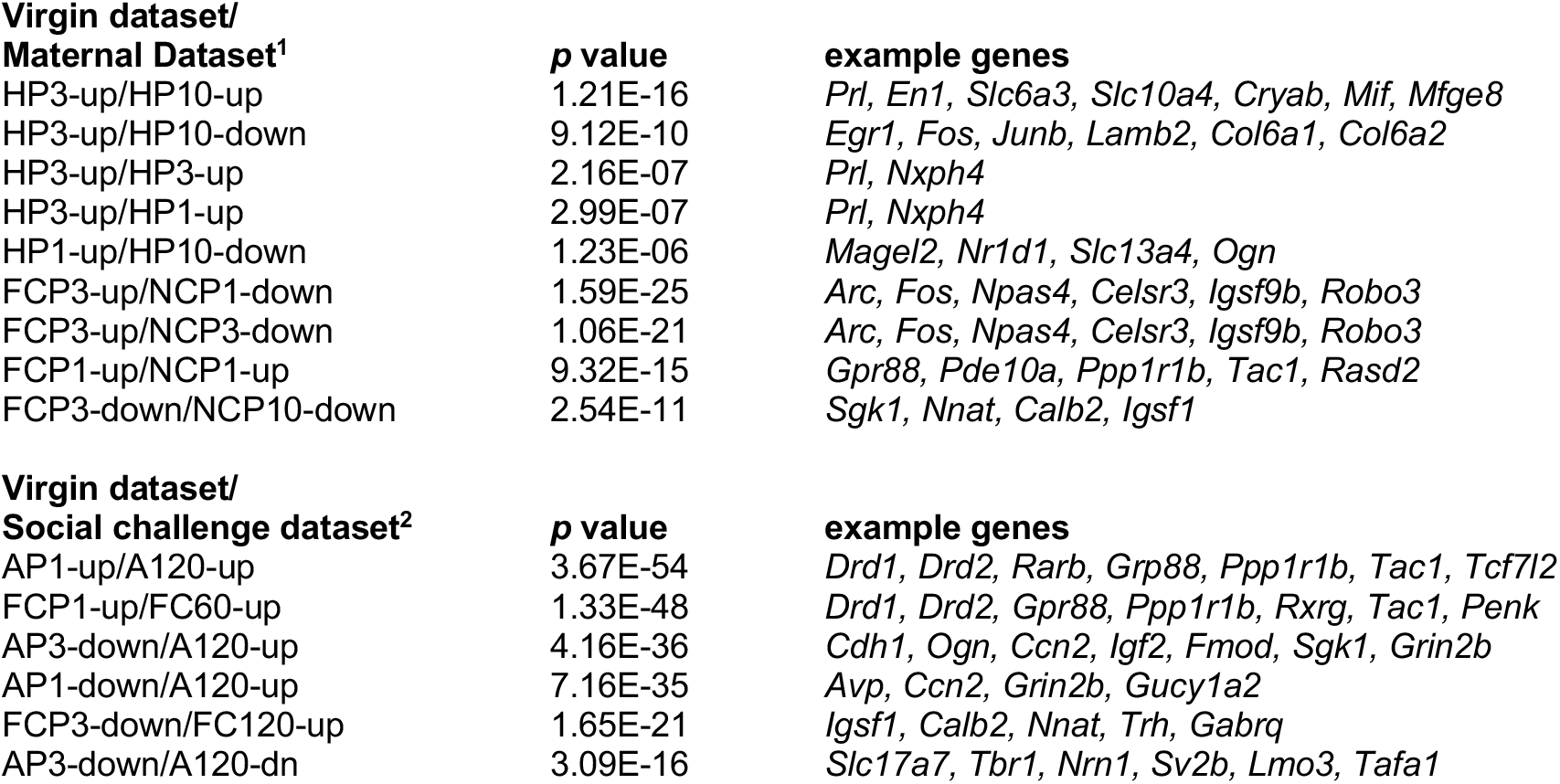
Significant correlations between differential gene expression in specific brain regions of pup-exposed virgins and new mothers, or virgins and socially challenged male mice. Hypothalamus (H), Amygdala (A), Frontal cortex (FC) and Neocortex (NC). Highest correlations for all comparisons involving at least 3 overlapping genes are shown, for a full list of comparisons see Table S2.

The overlapping genes included several involved in DA neuron development and function (*En1*, *Slc6a3* and *Slc10a4*), and neuroprotection and neuroinflammatory processes (*Cryab*, *Mif*, and *Mfge8*). On the other hand, *up-regulated* hypothalamic DEGs from P3 virgins also overlapped with genes that were *down-regulate*d in the maternal hypothalamus at P10 (**Table 1**); IEGs (*Egr1*, *Fos*, and *Junb*) dominated this list along with genes encoding extracellular matrix (ECM) proteins.

We further identified both positively and negatively-correlated overlaps in comparisons between virgin frontal cortex and maternal neocortex. DEGs *up-regulated* in virgin frontal cortex at P3 overlapped significantly with DEGs *down-regulated* in neocortex of mothers at P1 and P3 (**Table 1**); as in hypothalamus, this group of oppositely regulated genes was dominated by IEGs (*Npas4*, *Arc*) and genes involved in ECM, and more particularly ECM proteins involved in axon pathfinding (*Celsr3*, *Igsf9b*, *Robo3*). Interestingly, there was also significant overlap between DEGs *up-regulated* in virgin P1 frontal cortex and P1 maternal neocortex. This cluster included genes related to the anxiety-related response that, as noted above, were also up-regulated in the virgin P1 amygdala (*Adora2a*, *Gpr88*, *Pde10a*, *Rasd2*, *Ppp1r1b*, *Tac1*, *Syndigl1*) (**Fig. 3; Table S2B, S2C**); this finding suggested the possibility that mothers might also experience a similar anxiety-related reaction soon after pups were born.

More generally, DEGs across the maternal brain showed enrichment in many of the same functional categories detected in brains of the alloparenting virgin mice [39]. Although direct comparison of the same brain regions at similar time points will be required for further clarification, the data are consistent with the idea that virgin and maternal brains activate many of the same pathways in response to pups. We note the exception of activation of IEGs and plasticity-related ECM genes to this general pattern.

#### Gene expression in P1 virgins closely parallels that of socially challenged males

The similarities between gene expression in mothers and the P3 virgins fits well with the fact that the virgins were beginning to exhibit maternal behaviors around this time. However, the molecular events in the virgin frontal cortex and amygdala around P1 remained something of a puzzle. We noted some similarities between DEGs in the virgin P1 amygdala and frontal cortex and DEGs previously identified in the same brain regions taken from of male mice undergoing a territory threat [40], and a hypergeometric test confirmed a very robust correlation (**Table 1; Table S2D**). DEGs up-regulated in P1 virgin amygdala - and particularly those down-regulated in P3vP1 comparisons - showed especially high levels of overlap with genes up-regulated in the amygdala of the socially challenged males; frontal cortex DEGs followed a similar pattern. The overlapping amygdala genes included those associated with dopamine and cholinergic signaling (e.g. *Drd1*, *Drd2*) as well as a large cohort of TF-encoding genes (e.g. *Rarb*, *Foxp1*, *Neurod2*, *Tcf7l2*) (**Table S2E**). The common up-regulation of these genes in the two social contexts suggests an especially important and common role. The data are consistent with the interpretation that at P1, the virgin females are experiencing emotions related to fear and threat, marked by a genomic response that is remarkably similar to that operating in the brains of males involved in territory defense. Notably, this threat-related P1 transcriptomic response was largely extinguished in the virgins at P3, as the females began to display maternal behavior toward the pups.

### DEGs cluster into network modules, suggesting regulatory factors with coordinated roles

To gain insights into the coordination and interactions of regulatory factors involved in these brain transcriptomic events, we used a weighted gene correlation network analysis (WGCNA) approach [41] to generate a co-expression network, including 25 co-expression modules connected by positive or negative links (**Fig. 4A**, **Table S4A-D).** DEGs from particular brain regions and time points clustered strongly within certain network modules, indicating the coordinated regulation of functionally inter-related genes **(Table S4E**). In particular, the threat-related genes that were up-regulated in the virgin amygdala at P1 and then down-regulated at P3 compared to P1, were especially highly enriched in module 3, with modules 7 and 8 showing a similar but less robust enrichment pattern (**Fig. 4B**). These modules included most of the genes that were similarly expressed in P1 virgins and socially challenged males (**Table S2D**). The three positively correlated modules also included several sets of known interacting genes and DEGs with related functions. For example, Module 3 includes *Drd1* and *Penk* together with TF genes *Rarb* and *Foxp1*, both of which are important to development and activity of development of dopaminergic neurons [42,43]. Module 7 includes *Drd5* and TF-encoding DEG *Tcf7l2*, which has been implicated in fear learning [44]; Module 8 includes *Drd2*, *Pdyn*, *Tac1*, and *Rxrg*, the latter encoding RARB dimerization partner, RXRG. Therefore, the DEGs cluster into modules with inter-related functions, including TFs with known regulatory interactions.

**Figure 4.**
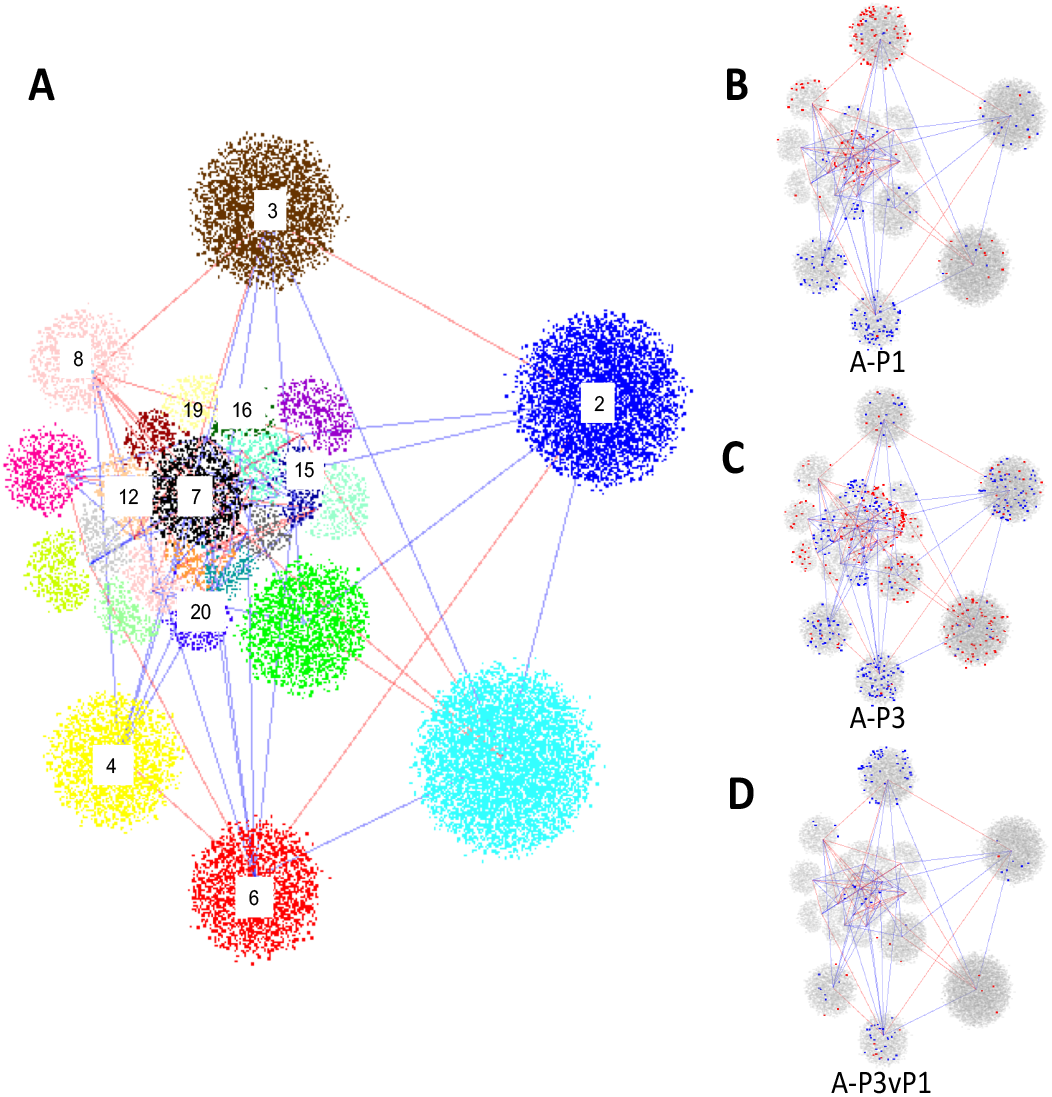
Weighted gene correlation network representing gene expression in four brain regions of virgin female mice. (A) The network, composed of genes expressed in Amygdala, Frontal Cortex, Hypothalamus, and Striatum of pup-exposed and non-exposed virgin controls, consists of 25 modules, each represented by clusters of different color and joined by lines representing positive (blue) or negative (red) eigengene correlations (in both cases showing only those correlations ≥0.6). Numbers have been added to label modules with particular enrichments in amygdala (A) DEGs as referred to in the text. (B-D) representations of the same network but showing the module membership of up-(blue dots) or down-regulated (red dots) DEGs identified in P1 v E18 (A-P1, B), P3 v E18 (A-P3, C), or P3vP1 (A-P3vP1, D) transcriptomic comparisons. Full details of network structure, membership and correlations are provided in Table S4.

Other DEG classes clustered into distinct network modules. For example, genes down-regulated at both P1 and P3 clustered together, especially in Modules 4 and 6 (**Table S4E**); P3 up-regulated genes clustered with especially high concentration in Module 15, including the heat shock factor regulator, *Hsf1*, a neuroprotective factor involved in adaptation to stressful experience [45]. Other modules displaying more modest levels of amygdala DEG enrichment reflect brain expression patterns that are strongly correlated with, or anticorrelated to, Modules 3, 6 or 15 and might thus also include regulatory factors involved in cross-module gene activation or repressive effects (**Table S4E**).

To identify TFs most central to the pup response, we used GENIE3 [46] to reconstruct a gene regulatory network (GRN) with these same data (Table S5A). We then identified TFs in the network with target gene sets that were most highly enriched in DEGs from each brain region and time point (**Table S5B-E**). The data pointed clearly to Module 3 TF *Rarb* as the most central TF in the amygdala P1 transcriptomic response, whereas Module 6 TF genes *Foxc2*, *Osr1*, and *Prdm6*, all three of which are down-regulated in P3 amygdala, dominated the amygdala P3 transcriptomic response (Table S5B); brain functions of these Module 6 TFs are not known. Module 15 TF gene *Hsf1*, which is itself up-regulated in hypothalamus at P3, was the most highly associated with DEGs in that brain region and time point (Table S5C). Of potential interest, *Snapc4*, a module 15 TF that activates expression of small nuclear RNAs [47,48] figured prominently in hypothalamus at both time points, suggesting a role for regulation of RNA splicing in the hypothalamic response.

### A dramatic shift in chromatin landscape during the long-term nurturance experience

#### Dynamic changes in amygdala chromatin at the P3 time point

The behavioral adaptations that follow maternal and alloparenting experiences have long been thought to involve epigenetic factors [23,26]. We therefore expected that histone modifications could play a key role in the virgins’ transition to maternal care. In particular, we hypothesized that the key genes involved in the threat reaction we observed at P1 might be actively silenced by these mechanisms as the virgins began to display maternal behaviors at P3. We tested this hypothesis by carrying out chromatin immunoprecipitation (ChIP) in chromatin from each of the four brain regions from virgin females co-housed with mothers at E18 and P3. For these ChIP experiments, we used an antibody specific to histone 3 acetylated at lysine 27 (H3K27Ac), a general marker for accessible chromatin [49]).

Consistent with our previous results [40], the ChIP profiles revealed tens of thousands of open-chromatin peaks in every brain region for both pup-exposed and non-exposed females (**Table S6**). Since differentially accessible peaks (DAPs) offer a unique window into chromatin dynamics that may drive the brain response, we paid special attention to DAP regions – defined as genomic regions in which the relative levels of H3K27Ac were at least two-fold higher or lower in brains collected at P3 compared to E18 consistently in biological replicate samples at FDR < 0.05 (**Table S7**). Surprisingly although peaks were identified in similar numbers overall in the each of the four brain regions, DAPs were virtually absent in the chromatin samples from hypothalamus at P3 and were found in relatively low numbers in frontal cortex and striatum at this time point as well. In striking contrast, chromatin from the P3 amygdala contained thousands of DAPs, either increased (5325 DAPs) or decreased in accessibility (7209 DAPs) at P3 compared to E18 (**Fig. 5A, Table S7A**). To maximize the chances of linking DAPs to specific DEGs, we focused our attention on the smaller number of DAPs located within 5 kb of the TSS of an annotated gene (called TSS-DAPs). Altogether we found 2738 TSS-DAPs with decreased H3K27Ac at P3 compared to E18 abbreviated hereafter as P3-closed DAPs), and 1040 TSS-DAPs with increased levels of H3K27Ac accumulation at P3 compared to E18 (P3-open DAPs). Interestingly, all genes associated with P3-open or P3-closed TSS-DAPs, respectively, clustered into network modules that were also enriched for up-or down-regulated DEGs. For example, amygdala P3-closed DAPs were particularly enriched for linkage genes in modules 3 and 6, whereas P3-open DAPs were most likely to be associated with genes in Modules 7 and 8 (**Table S4F**). The TSS-DAPs were associated with 138 amygdala DEGs, including 22 TSS containing P3-open DAPs, 115 TSS containing P3-closed DAPs, and 1 TSS hosting DAPs of both types (**Table S6B**). Given that amygdala DEGs were up-and down-regulated in roughly equal numbers (Fig. 2) the preponderance of down-regulated genes in TSS-DAPs suggested that alterations in the chromatin landscape at P3 were primarily focused on silencing amygdala genes. The DEGs associated with P3-closed DAPs were enriched specifically and significantly in network module 3 (hypergeometric *p* = 2.57E-17), suggesting an especially important role for histone de-acetylation in silencing this coregulated cluster of threat-associated genes.

**Figure 5.**
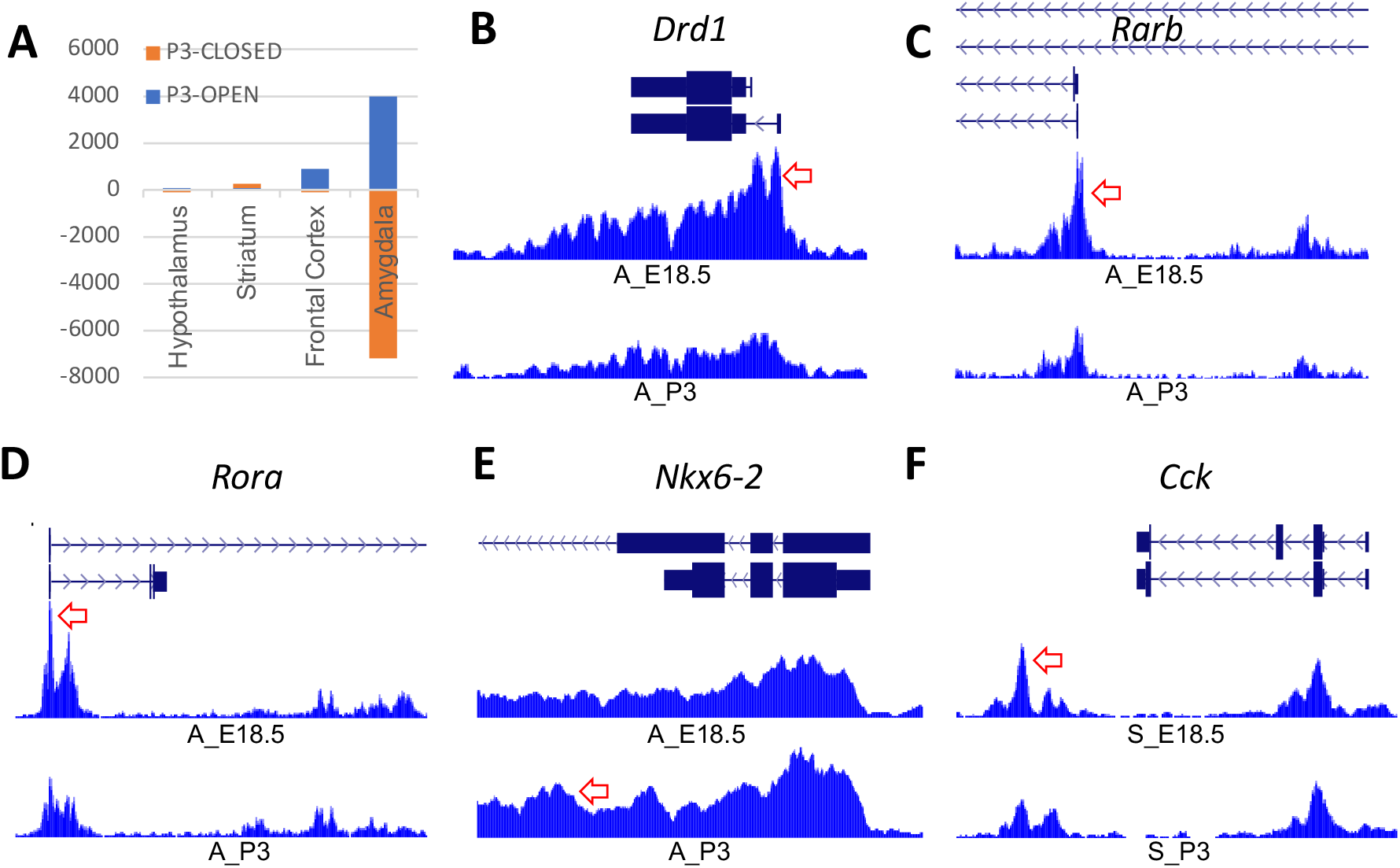
Dramatic changes in the amygdala chromatin landscape accompanies the transition to maternal-like behaviors in alloparenting virgin mice. (A) Relative numbers of TSS-associated differentially accessible peaks (DAPs), as measured by >2-fold change in detected levels of H3K27Ac, in the four brain regions tested in this study. Positive numbers represent peaks more accessible at P3 than E18 (P3-open DAPs); negative numbers represent less accessible (P3-closed) peaks. (B-E) Examples of DAPs in *Drd1*, *Rarb1*, *Rora*, *and Nkx6-2* genes, showing normalized H3K27Ac profiles in amygdala chromatin at E18 (A_E18, top track) and P3 (A_P3, bottom track) for each gene. (F) An example of a DAP in chromatin from Striatum (S_E18, S_P3) within the *Cck* gene. Red arrows in each panel point to examples of significant differentially accessible peaks. Full ChIP profiles are available online as a UCSC Browser track hub at https://trackhub.pnri.org/stubbs/ucsc/public/allo.html), and data are available in Tables S5 and S6.

Examples include amygdala P3-closed DAPs associated with TSS of Module 3 genes, *Drd1* and *Rarb* (**Fig 4B, 3C**), P3-closed DAPs associated with the primary alternative promoter of *Rora* (Module 6 TF gene down-regulated at both P1 and P3) (**Fig. 4D**), and P3-open DAPs associated with *Nkx6-2* (Module 1 TF gene up-regulated at the P3 timepoint) (**Fig. 4E**). Together the data suggest that activities of many key genes involved in pup response are regulated by differential chromatin accessibility, specifically in the amygdala.

Although much smaller in numbers, some DAPs associated with DEGs in other brain regions also deserve some mention. For example, *Rarb* was also down-regulated in frontal cortex P3 vs. P1 comparisons, and displayed a pattern of DAPs in cortex very similar to that seen in amygdala (**Table S6A**). Additional DAPs that may be relevant to gene expression were discovered in brain regions other than amygdala by lowering the fold-change cutoff to 1.5 instead of 2 (1.5X vs 2X), while keeping the same replicate FDR significance threshold (FDR < 0.05). For example, a 2X P3-open DAP in striatal chromatin was identified approximately 25 Kb downstream of *Cck*, and several 1.5X P3-closed DAPs were found within and closer to the gene (**Table S6A; Fig. 4F)**. Since *Cck* was up-regulated in striatum at P3, this chromatin configuration suggests the possible role for chromatin dynamics in the regulation of this critical gene.

#### Enrichment of binding motifs points to mechanistic insights

To obtain further information regarding the potential activity of TFs in the pup response, we searched for enrichment of known TF binding motifs (TFBMs) in the P3-open and P3-closed amygdala TSS DAPs. The search identified REST/NRSF binding motifs as the top enrichment within *P3 closed* TSS-DAPs (E18 enriched compared to P3); although *Rest* itself was not identified as differentially expressed, the data suggest that REST TFBMs were being closed between E18 and P3 in amygdala of the pup-exposed virgins. This finding is of interest, because REST is a central regulatory of neuron differentiation and plasticity [50], and also plays a role in stress resilience in adult brain [51]. The search also identified enrichments for motifs in the P3-closed DAPs that are recognized by TFs encoded by amygdala DEGs, including P3 down-regulated genes *Mef2c* and *Rora* (Table S2). Consistent with their expression, the TSS of both genes were associated with P3-closed DAPs (**Fig. 4D; Table S7B**). Together these data indicate that histone deacetylation events evident at P3 serve to not only reduce levels of *Mef2c* and *Rora* gene expression, but simultaneously, to reduce the accessibility of both TFs to their target genes. Notably, both TF genes have been associated with deficits in social behavior [52–54] and *Rora* has been implicated in maternal behavior specifically [55], supporting a functional role. Notably, *Mef2c* target genes predicted by GRN analysis were significantly and specifically enriched in pup-driven DEGs in amygdala; MEF2C target genes were also predicted to include a notably high number of other TFs (Table S5B). These data suggest that MEF2C may play a role as one of the central hubs coordinating the amygdala transcriptomic response.

Some notable TFBM enrichments were also detected in P3-open DAPs. For example, we noted enrichment of FOX family TFs including the specific TFBM of the protein encoded by DEG, *Foxp1* which was up-regulated in amygdala at P1, then down-regulated between P1 and P3 (**Table S7**). However, this FOX motif could potentially also be recognized by other family members including FOXC2; *Foxc2* was down-regulated in the P3 amygdala and was predicted in the GRN to be central to the P3 response (Table 3). Because we did not measure chromatin at P1, the initial timing of these epigenetic events is not discernible. However, the data suggest that while TSS-linked targets of REST, RORA and MEF2C became less accessible, targets of FOX family proteins became more accessible via epigenetic modifications during the postnatal period.

## DISCUSSION

With the goal of understanding molecular mechanisms that underlie the transition to maternal behaviors, we investigated the behavioral, transcriptomic, and epigenomic response of virgin females as they were co-housed with mothers and newborn pups over a period of several days. A recent study used this same paradigm to show that alloparenting virgins are instructed in pup care by co-housed mothers [12], and we observed a very similar pattern of behaviors in the mothers and virgins we tested here. Along the way, although virgin female mice did not display an obvious aversion toward the pups, we also observed evidence of an initial antagonism during the first two postnatal days; these antagonistic behaviors gave way to increasing levels of attention to the pups, with the virgins increasingly licking, grooming and huddling over the pups by postanatal day 3.

The data presented here reveal a dramatic and dynamic neurogenomic shift that coincides with successful maternal instruction and the activation of oxytocin neurons in the virgin brain. In particular, at P1 we observed a striking signal of fear and anxiety in the virgin hypothalamus, frontal cortex and especially in the amygdala, in the form of a gene expression pattern that correlated with high significance to that observed in territory-challenged males [40]. Despite the lack of obvious aversion, these data indicate that indeed - at least within this co-housed paradigm - virgin female mice do initially perceive the pups as anxiety-inducing, or even possibly threatening. If the threat signal is related to the aversive response observed in rats and other species, the data would be consistent with the hypothesis that pup aversion and defensive behaviors share a common brain circuitry [4], and would suggest a shared molecular mechanism for diverse types of threat response as well. Several TF genes implicated by gene expression, network co-expression, chromatin analysis, and/or motif-enrichment analysis were similarly up- or down regulated in the pup-exposed virgins and socially challenged males, suggesting crucial roles for these TFs in regulating this shared molecular signature of social threat.

Published data support the roles of several of these TFs in threat/anxiety response. For example, the RARB:RXRG dimer’s activities in amygdala have been linked to expression of anxiety-related phenotypes [56], and *RORA* is associated with enhanced fear response in humans [57] and mice [58]. Furthermore, *Rora* mutant mouse mothers do not retrieve, care for, or suckle their young [59]; the data presented here support further investigation of this gene’s role in maternal amygdala. Furthermore, other TF genes implicated in the shared threat signature are associated with social-behavior phenotypes, including *Tcf7l2*, which is up-regulated in the amygdala of virgins at P1 as well as in socially challenged males [40] and is important for fear-learning and adaptation [44]. Our data suggest that these TFs have coordinated roles in the fear response, with *Rarb* playing a central role.

We hypothesize that these and other networked TFs work together to modulate the response to pups, and the networks developed from our dataset suggests a robust framework of positive and negative gene interactions that coordinate this behavioral switch over time. The expression of the threat-related TF genes was extinguished along with the pulse of dopamine signaling after the rise of oxytocin, prolactin, and other neuropeptides by P3, paving the way for a shift in the virgin females’ behavior toward the pups; we surmise that this shift was driven, at least in part, by a substantial level of chromatin remodeling in the amygdala. Many of the genes that returned to normal expression levels between P1 and P3 in amygdala were associated with differentially accessible chromatin, consistent with their active epigenetic silencing in that brain region at P3.

Chromatin remodeling has been implicated in the development of maternal behaviors in both mothers and alloparenting virgins, although most published studies have focused on the hypothalamic MPOA as the primary site of this epigenetic response [16,26,60]. These studies have shown that *suppression* of HDAC activity – and thus *inhibition* of chromatin silencing - in hypothalamus is key to driving the females’ maternal response. Surprisingly therefore, we found no evidence of chromatin remodeling in the P3 hypothalamus, and the massive chromatin remodeling we did observe in amygdala was weighted toward *de-acetylation*, or chromatin closure, along with the silencing of differentially expressed genes. These findings would suggest that HDAC activity plays a positive role in the acquisition of maternal behavior, although it is certainly possible that histone deacetylation at earlier time points or in different brain regions could have been crucial. For example, although deacetylation in amygdala may be critical in quenching the threat response once established, *suppression* of deacetylation in hypothalamus (and/or other brain regions) before P1 might have prevented the establishment of the aversive/fear response in the first place. Possibly relevant to this hypothesis is the coordinated up-regulation of histone deacetylase and chromatin remodeling genes we detected in the hypothalamus at P3; this signal could reflect the trace of earlier epigenetic events associated with the expression of fear and anxiety in the virgins before they transitioned to active pup care.

This is the first study to investigate global gene expression in the amygdala in the context of alloparental care, and supports the idea of including amygdala in future studies with mothers as well. Our studies highlight a special role for the amygdala in the switch to alloparenting behavior in this context, a hypothesis that is consistent with the known functions of amygdala in maternal behavior and bonding [61,62]. As underscored by human brain imaging studies, maternal behavior involves a global brain response that unfolds over an extended period of pre- and postnatal time [63]; in the pup-exposed virgins, we detected just the start of this behavioral transition during the third postnatal day. Nevertheless, the mechanisms involved in this transition to intensive pup care could be relevant to a successful transition to motherhood as well. Although it is not yet possible to determine whether a similar response is activated, or actively suppressed, at some time around birth in the maternal amygdala, this question is an important one in the context of maternal bonding and infant care. Especially given the similar up-regulation of anxiety-related genes we identified in published data from the maternal neocortex, we speculate that a similar active suppression of a threat/anxiety program may occur in the amygdala of new mothers, and that dysregulation of this program could underlie the failure of mother-infant bonding, post-partum anxiety and depression. Addressing this hypothesis will offer a novel perspective on the causes of these very common, painful and highly consequential human maladies.

## MATERIALS AND METHODS

### Mice and behavioral analysis

All work with mice was done under the approval of the IACUC at the University of Illinois, Urbana Champaign. Mice were housed in a temperature-controlled room in a reverse 12h/ 12h light-dark cycle. Six-week-old female mice (C57BL/6J x C3HJ F1 hybrids, with an agouti coat to allow clear distinction with the black-coated virgins) purchased from the Jackson Laboratory were impregnated, and co-housed with age-matched virgin female C57BL/6J mice during pregnancy and through the early post-partum pregnancy. To record behavior, four pairs were filmed in clear-topped cages continuously using a Samsung SCB-2000 CCTV Camera with iSpy 64 v7.2.1.0 CCTV software. Behavior was scored in each cage (0 or 1) in 5-minute snapshots at the top of each hour from the day before and until the end of the fourth day after birth, with scores combined over 6 h periods for each cage to generate the illustrative plot in Fig. 1. To test the hypothesis that pup-grooming behaviors increased over time, while mother-grooming behaviors did not, we performed one-way repeated measures ANOVA in R (v4.0.4) using the rstatix package (v0.7.0). The anova_test() function in rstatix automatically assesses repeated measures data for the assumption of sphericity. Scores for these and other behaviors are also presented in Table S1. The 5-minute video snapshots are provided as File S1 with video clips of specific and unusual behaviors noted in the text provided as File S2; additional video is available on request.

### Dissections and RNA preparation

Dissections were performed as described in detail previously [40] with the addition of the striatum. Briefly, mice were euthanized by cervical dislocation followed by rapid decapitation. Their brains were removed and sectioned in a coronal slicing mouse brain matrix. A total of three cuts were made: two cuts separated by 4 mm defined by the rostral and caudal aspects of the hypothalamus and a third cut bisecting these two cuts. The hypothalamus, frontal cortex, striatum, and amygdala were dissected from the resultant brain slices (**Supplementary Figure S1**). Upon completion of these dissections, focal brain regions were placed into cryotubes, snap-frozen in liquid N2, and stored at −80°C until downstream processing. Samples were prepared for sequencing from the four dissected brain regions of five mice per condition (E18, P1, P3). RNA isolation and QC were completed as described previously, with libraries prepared robotically at the Roy J. Carver Biotechnology Center at University of Illinois, also as described in [40].

### Gene expression analysis

Illumina sequencing libraries were generated with the TruSeq Stranded mRNA HT kit (Illumina) using an Eppindorf ePMotion 5075 robot and were sequenced to a depth 45-60 million reads per sample on Hi-Seq 2500 instruments at the Roy J. Carver Biotechnology Center at the University of Illinois. All sequencing data generated in this study have been deposited to the GEO database under Accession number GSE184549. Pairwise comparisons of E18, P1 and P3 samples were completed as previously described in detail [40], with results provided in Table S2. For functional analysis, genes that were found to be differentially expressed at fdr < 0.05 were first filtered for absolute fold change >1.5, and uploaded to the ToppCluster web analysis tool [64] using default conditions (Bonferroni correction, fdr <0.05). Selected categories are summarized in Table 2, with full ToppCluster Results reported in Table S3.

### Network Analysis

We used signed WGCNA (Langfelder & Horvath, 2008) to generate networks from the data from all individuals, brain regions, and time points, as described in depth in our previous study [40]. Eigengenes calculated for each module were used to generate module correlations; details of module structure, module gene content, eigengene correlations, and hypergeometric enrichments are presented in Table S4. After log-transforming our data using voom+limma, we filtered zero variance genes, selected a soft thresholding coefficient of 3, then used a signed Pearson correlation analysis with a minimum module size of 30. Images in Figure 3 were generated using version 3.7.1 of Cytoscape [65].

To reconstruct the GRN, we obtained a list of 1523 potential transcription factors in mouse from Animal Transcription Factor Database [66]. GENIE3 [46] was applied on the expression data of 37991 genes in 53 conditions consisting of various brain regions (H=hypothalamus, A=amygdala, FC=frontal cortex, and S=striatum) and different time points (E18, P1-P3) to score the relative significance of each TF-gene interaction. (Auto-regulatory relationships were excluded). To construct a GRN, for each gene we collected up to top five TF regulators of that gene as predicted by GENIE3, additionally requiring that the TF-gene pair have a Spearman’s correlation of at least 0.5 (in absolute value) and a GENIE3 score of at least 0.005. The resulting GRN included 92717 interactions involving 1400 unique TFs and 21156 genes (Table S5A). To assess the significance of TF regulons in different brain regions, for each TF we computed the enrichment of its regulon (gene set predicted to be regulated by the TF) for DEGs from each brain region, using hypergeometric test (Table S5B-E).

### ChIP Tissue Preparation, Chromatin Immunoprecipitation, and Library Preparation

ChIP was performed essentially as described in detail in our previous study [40]. Briefly, brain tissue dissected from 3 animals was pooled, homogenized, and fixed in PBS with 1% formaldehyde for 10 minutes. Nuclei were prepared from the fixed cells and stored at −80° C until use. Thawed nuclei were sonicated using a BiorupterTM UCD-200 (Diagenode, Liège, Belgium) sonicator, and fragmented chromatin was processed for ChIP with 2 ug histone H3K27Ac antibody per sample (Abcam ab4729), using one million nuclei for each IP. IPs were performed in biological replicate, with one pool of 3 samples in each replicate, as previously described. Libraries were prepared from eluted DNA using KAPA LTP library kits (KK8230) using Bioo Scientific index adapters, size-selected using AmpureXP beads (Beckman Coulter, Brea, CA, USA) and quality checked by Qubit 2.0 and Bioanalyzer (Agilent 2100). Samples were sequenced to a depth of 20-30M reads per replicate on an Illumina HiSeq 2500 sequencer using a TruSeq SBS sequencing kit, version 4, in single-end format with fragment length of 100 bp. Base calling and demultiplexing into FASTQ files was done using bcl2fastq v1.8.4 software (Illumina, San Diego, CA, USA).

### ChIP-Seq Bioinformatics

ChIP sequencing reads were mapped with Bowtie2 [67] to the UCSC Mus musculus mm9 or mm10 genome, using default settings and analyzed for peaks using HOMER (Hypergeometric Optimization of Motif EnRichment) v4.7 [68], as previously described [40]. Differential chromatin peaks were identified in biological replicates using the HOMER getDifferentialPeaksReplicates.pl script, looking for any peaks that changed at least two-fold between conditions with an FDR cutoff of 0.05. Known motif discovery was performed with the HOMER findMotifsGenome.pl script using default settings with 201bp peak regions extracted from all histone peaks or only differential histone peaks. Chromatin profiles are available online as a UCSC Genome Browser track hub at https://trackhub.pnri.org/stubbs/ucsc/public/allo.txt).

## ACKNOWLEGEMENT

We thank Gene Robinson, Alison Bell, Dave Zhao for helpful discussions throughout this project and Alison Bell and Elbert Branscomb for critical comments on the manuscript. This work was supported by the Simons Foundation (#SfLife 291812), with additional funding provided by the University of Illinois and the Pacific Northwest Research Institute.

