## Supplementary figures and images for "An epigenomic shift in amygdala marks the transition to maternal behaviors in alloparenting virgin female mice" (from another version of this preprint: version 1, 2021-05-17)

### Fig. S1

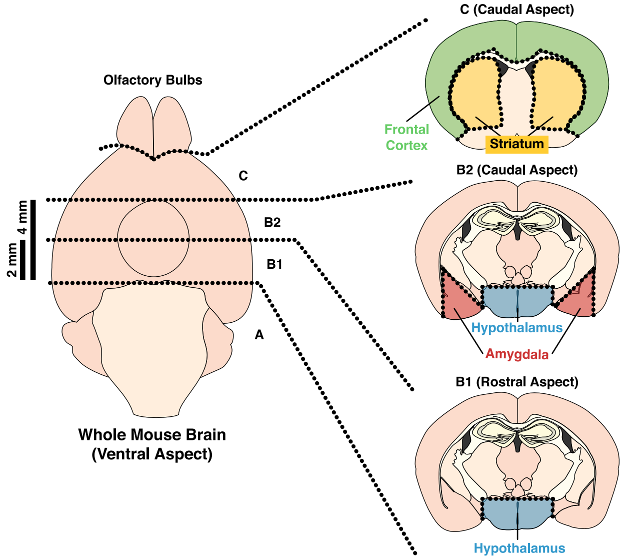
